## Supplemental Material for "Different coexistence patterns between apex carnivores and mesocarnivores based on temporal, spatial, and dietary niche partitioning analysis in Qilian Mountain National Park, China"

**Table S1** Relative frequencies of prey species consumed in carnivore diets (SL-snow leopard, EL-Eurasian lynx, PC-Pallas's cat, RF-Red Fox, TF-Tibetan Fox. 'N' represents the number of scats).

| | | Overall | Wolf | Snow Leopard | Eurasian Lynx | Pallas's cat | Red Fox | Tibetan Fox |
|---|---|---|---|---|---|---|---|---|
| | | N=404 | N=49 | N=147 | N=19 | N=63 | N=87 | N=39 |
| Livestock | Domestic Goat | 0.18% | 1.45% | 0.00% | 0.00% | 0.00% | 0.00% | 0.00% |
| | Domestic Sheep | 1.65% | 5.80% | 0.92% | 0.00% | 0.00% | 2.50% | 0.00% |
| | Domestic Yak | 10.64% | 17.39% | 13.76% | 26.67% | 0.00% | 5.83% | 2.33% |
| Artiodactyla | Blue Sheep | 17.98% | 21.74% | 31.65% | 0.00% | 1.54% | 10.00% | 2.33% |
| | Wild Yak | 4.22% | 0.00% | 7.80% | 0.00% | 0.00% | 5.00% | 0.00% |
| | Musk Deer | 0.55% | 0.00% | 1.38% | 0.00% | 0.00% | 0.00% | 0.00% |
| Carnivora | Eurasian Badger | 0.18% | 0.00% | 0.46% | 0.00% | 0.00% | 0.00% | 0.00% |
| | Pallas's Cat | 0.37% | 1.45% | 0.46% | 0.00% | 0.00% | 0.00% | 0.00% |
| | Red Fox | 0.18% | 1.45% | 0.00% | 0.00% | 0.00% | 0.00% | 0.00% |
| | Tibetan Fox | 0.37% | 1.45% | 0.46% | 0.00% | 0.00% | 0.00% | 0.00% |
| Lagomorpha | Plateau Pika | 38.35% | 27.54% | 14.68% | 16.67% | 92.31% | 46.67% | 86.05% |
| | Woolly Hare | 5.50% | 4.35% | 5.05% | 30.00% | 1.54% | 5.00% | 0.00% |
| Rodentia | Chinese Birch Mouse | 0.55% | 0.00% | 0.00% | 3.33% | 0.00% | 0.83% | 2.33% |
| | Irene's Mountain Vole | 3.30% | 0.00% | 1.83% | 13.33% | 1.54% | 7.50% | 0.00% |
| | Himalayan Marmot | 11.38% | 14.49% | 19.27% | 6.67% | 0.00% | 5.00% | 4.65% |
| | *Eospalax sp.* | 1.10% | 0.00% | 0.00% | 3.33% | 0.00% | 4.17% | 0.00% |
| | House Mouse | 0.18% | 0.00% | 0.00% | 0.00% | 0.00% | 0.00% | 2.33% |
| Galliformes | Himalayan Snowcock | 0.37% | 1.45% | 0.46% | 0.00% | 0.00% | 0.00% | 0.00% |
| | Tibetan Partridge | 0.37% | 0.00% | 0.00% | 0.00% | 0.00% | 1.67% | 0.00% |
| Passeriformes | *Montifringilla sp.* | 0.18% | 0.00% | 0.00% | 0.00% | 0.00% | 0.83% | 0.00% |
| | *Prunella sp.* | 0.18% | 0.00% | 0.00% | 0.00% | 0.00% | 0.83% | 0.00% |
| | Hume's Groundpecker | 0.18% | 0.00% | 0.00% | 0.00% | 0.00% | 0.83% | 0.00% |
| Columbiformes | Eurasian Collared Dove | 0.18% | 0.00% | 0.00% | 0.00% | 1.54% | 0.00% | 0.00% |
| Strigiformes | Eurasian Eagle Owl | 0.18% | 0.00% | 0.46% | 0.00% | 0.00% | 0.00% | 0.00% |
| Falconiformes | *Buteo sp.* | 1.47% | 1.45% | 1.38% | 0.00% | 1.54% | 2.50% | 0.00% |
| | *Falco sp.* | 0.18% | 0.00% | 0.00% | 0.00% | 0.00% | 0.83% | 0.00% |

**Table S2** Dietary diversity for each carnivore species. The higher the value, the greater dietary diversity observed.

| | Richness | Shannon-Wiener Index |
|---|---|---|
| Wolf | 12 | 1.941 |
| Snow Leopard | 15 | 1.943 |
| Eurasian Lynx | 7 | 1.688 |
| Pallas's Cat | 6 | 0.395 |
| Red Fox | 16 | 1.980 |
| Tibetan Fox | 6 | 0.622 |

**Table S3** Jaccard's distance of prey items in diets.

| | | Inversed Jaccard's index |
|---|---|---|
| Apex vs. Apex carnivores | Wolf – Snow leopard | 0.588 |
| | Wolf – Eurasian lynx | 0.267 |
| | Snow leopard – Eurasian lynx | 0.294 |
| Apex vs. Mesocarnivores | Wolf – Pallas's cat | 0.286 |
| | Wolf – Red fox | 0.333 |
| | Wolf – Tibetan Fox | 0.286 |
| | Snow leopard – Pallas's cat | 0.313 |
| | Snow leopard – Red fox | 0.409 |
| | Snow leopard – Tibetan Fox | 0.235 |
| | Eurasian lynx – Pallas's cat | 0.300 |
| | Eurasian lynx – Red fox | 0.438 |
| | Eurasian lynx – Tibetan Fox | 0.444 |
| Mesocarnivores vs. Mesocarnivores | Pallas's cat – Red fox | 0.294 |
| | Pallas's cat – Tibetan Fox | 0.200 |
| | Red fox – Tibetan Fox | 0.294 |

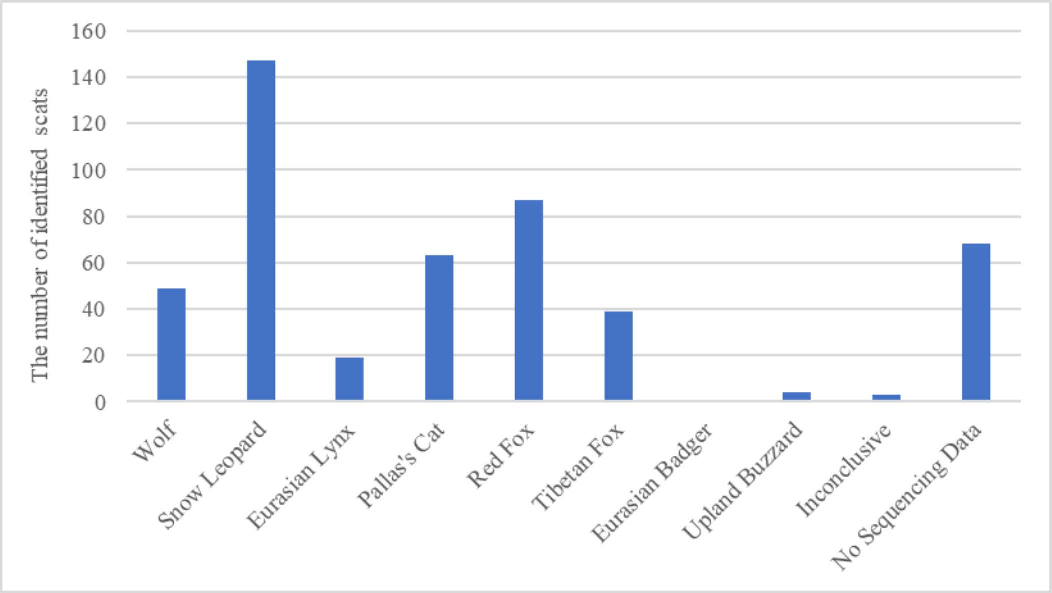


**Figure S1** The number of scats belonging to each host species among 480 scats samples.
